# Early infection response of the first trimester human placenta at single-cell scale

**DOI:** 10.1101/2023.01.02.522155

**Authors:** Regina Hoo, Elias R. Ruiz-Morales, Iva Kelava, Carmen Sancho-Serra, Cecilia Icoresi Mazzeo, Sara Chelaghma, Elizabeth Tuck, Alexander V. Predeus, David Fernandez-Antoran, Ross F. Waller, Marcus Lee, Roser Vento-Tormo

## Abstract

Placental infections are a major worldwide burden, particularly in developing countries. The placenta is a transient tissue located at the interface between the mother and the fetus. Some pathogens can access the placental barrier resulting in pathological transmission from mother to fetus, which may have a profound impact on the health of the developing fetus. Limited tissue accessibility, critical differences between humans and mice, and, until recently, lack of proper *in vitro* models, have hampered our understanding of the early placental response to pathogens. Here we use single-cell transcriptomics to describe the placental primary defence mechanisms against three pathogens that are known to cause fetal and maternal complications during pregnancy - *Plasmodium falciparum, Listeria monocytogenes* and *Toxoplasma gondii*. We optimise *ex vivo* placental explants of the first-trimester human placenta and show that trophoblasts (the epithelial-like cells of the placenta), and Hofbauer cells (placental macrophages) orchestrate a coordinated inflammatory response after 24 hours of infection. We show that hormone biosynthesis and transport are downregulated in the trophoblasts, suggesting that protective responses are promoted at the expense of decreasing other critical functions of the placenta, such as the endocrine production and the nourishment of the fetus. In addition, we pinpoint pathogen-specific effects in some placental lineages, including a strong mitochondrial alteration in the Hofbauer cells in response to *T. gondii*. Finally, we identify adaptive strategies and validate nutrient acquisition employed by the *P. falciparum* during placental malaria infection. This study provides the first detailed cellular map of the first-trimester placenta upon infection and describes the early events that may lead to fetal and placental disorders if left unchecked.

## Introduction

The placenta is a transient extra-embryonic organ located at the boundary between the mother and the fetus. During the nine months of human pregnancy, the placenta provides vital functions for the embryo including transport of nutrients and gases, hormonal production and protection of the fetus by acting as a selective barrier between the mother and fetus. Trophoblasts are the epithelial-like cells of the placenta that organise into villi structures floating in the uterine space. Villous cytotrophoblasts (VCTs) are the proliferative core of trophoblast that can fuse into multinucleated syncytiotrophoblast (SCT) which surrounds the placental villi. Some placental villi can anchor the endometrium, the mucosal layer of the uterus, and at this site, the extravillous trophoblasts (EVTs) detach from the placenta and infiltrate the endometrium^1–4^. Thus, the placenta establishes two distinct interfaces with the mother: (i) the intervillous space, where SCT are in direct contact with maternal blood, and (ii) the implantation site, where EVTs intermingle with maternal endometrial cells.

The placenta is a protective, efficient physical barrier that separates the maternal and fetal circulation. However, some pathogens, including parasites, bacteria and viruses can cross the placenta and enter the fetal circulation (i.e. vertical transmission)^5^. Placental infections can cause fetal defects, such as microcephaly^6^ and hydrocephalus^7, 8^, and lead to pregnancy complications, including stillbirth and fetal growth restriction^9^. Pathogens can enter the trophoblast barrier by hijacking existing receptors on the trophoblasts^10, 11^. The bacterium *Listeria monocytogenes* (*L. monocytogenes*) engages its internalin protein with placental E-cadherin and c-Met expressed on SCT to facilitate entry from the maternal environment^12, 13^. *Toxoplasma gondii* (*T. gondii*) is known to invade the trophoblast in order to maintain its intracellular replication lifestyle^14^ but the receptors used for entry are unknown. *Plasmodium falciparum* (*P. falciparum*) does not cross the placenta but can attach to it through the parasite’s variant PfEMP1 receptor VAR2CSA that binds to SCT proteoglycan chondroitin sulphate A (CSA)^15, 16^. Placental malaria, which is caused by *P. falciparum,* can cause adverse pregnancy outcomes including maternal anaemia, low birthweight and preterm delivery^16, 17^ and in severe cases, can lead to prenatal and maternal mortality^18, 19^. Despite the prevalence of placental malaria^17^ and other infections^20, 21^, particularly in low- and middle-income countries, as well as the severity of infections for both the mother and the fetus, the molecular mechanisms driving the infection and the interplay between pathogen virulence strategies and host defence strategies remains poorly studied in humans especially during the first-trimester of pregnancy.

The first-trimester embryo is particularly vulnerable to infections due to the adaptive immune response not being fully established^22^, and hence its protection relies on the innate responses in the placental barrier. Despite the relevance and impact of infection of the first-trimester placenta, this has been poorly studied^5^, mainly due to the obvious ethics and logistical limitations, and the restricted availability of *in vitro* models that account for the diversity of cell lineages present in the placenta. Moreover, an important hurdle is to extrapolate the protective and pathological mechanisms from mouse to humans due to critical differences in placental biology across species^23^. Trophoblast stem cells expanded *in vitro*^24^ and self-renewal primary trophoblast organoids^25, 26^ can recapitulate some aspects of placental development, and have recently been used to model Zika^27^ and human cytomegalovirus^28^ infection. However, other placental cell types which are likely to have an important role in the inflammatory responses in the placental barrier, including Hofbauer cells (fetal macrophages) and fibroblasts, are absent. An attractive *in vitro* model recently used to study infection in the uterine-placental interface^29, 30^ is the placental explant culture where small biopsy sections of the placental villous are cultured *in vitro*^31–33^. An accurate quantitative benchmark of these models at the single-cell level is required to describe the immune responses in the placenta *in vitro* and its utility to study early immune responses in this interface.

Single-cell and spatial transcriptomics methods have been transformative in understanding placental development^34, 35^, mapping the transcriptome of the parasites^36–38^ and studying host-pathogen interactions^39, 40^. Here, we optimise first-trimester placental explants and use them to study host-pathogens interactions at the single-cell level. We quantify the coordinated multilineage placental responses to *P. falciparum, T. gondii* and *L. monocytogenes* infection during the acute infection phase (24 hours). We identify early features of infection in the trophoblast and macrophage compartment that may promote placental disorders and provoke pathogenic transmission into the developing fetus if left unchecked. Finally, we demonstrate that malaria parasites undergo adaptation as they infect the human placenta. Thus, we provide a comprehensive study of the acute response of the human early placenta to pathogens, gaining mechanistic insights into host-pathogen interaction dynamics in this unique tissue microenvironment.

## Results

### Human placental explants recapitulate main lineages of the early placenta

We adapted the placental explant technique^33^ for *ex vivo* culture of primary placental tissue to quantify the response of the human placenta to acute infections by three pathogens (*P. falciparum, T. gondii* and *L. monocytogenes*) **(Figure 1a)**. In brief, fresh placental explants were dissected into 1 cm^2^ and grown on Matrigel-coated Transwells in modified trophoblast media (TOM)^25^ **(Supplementary Figure 1a-b)**. These culture conditions retain all the placental cell lineages and all cell types, including immune cells, and have high cell viability up to 72 hours in culture **(Supplementary Figure 2a-d)**.

**Figure 1.**
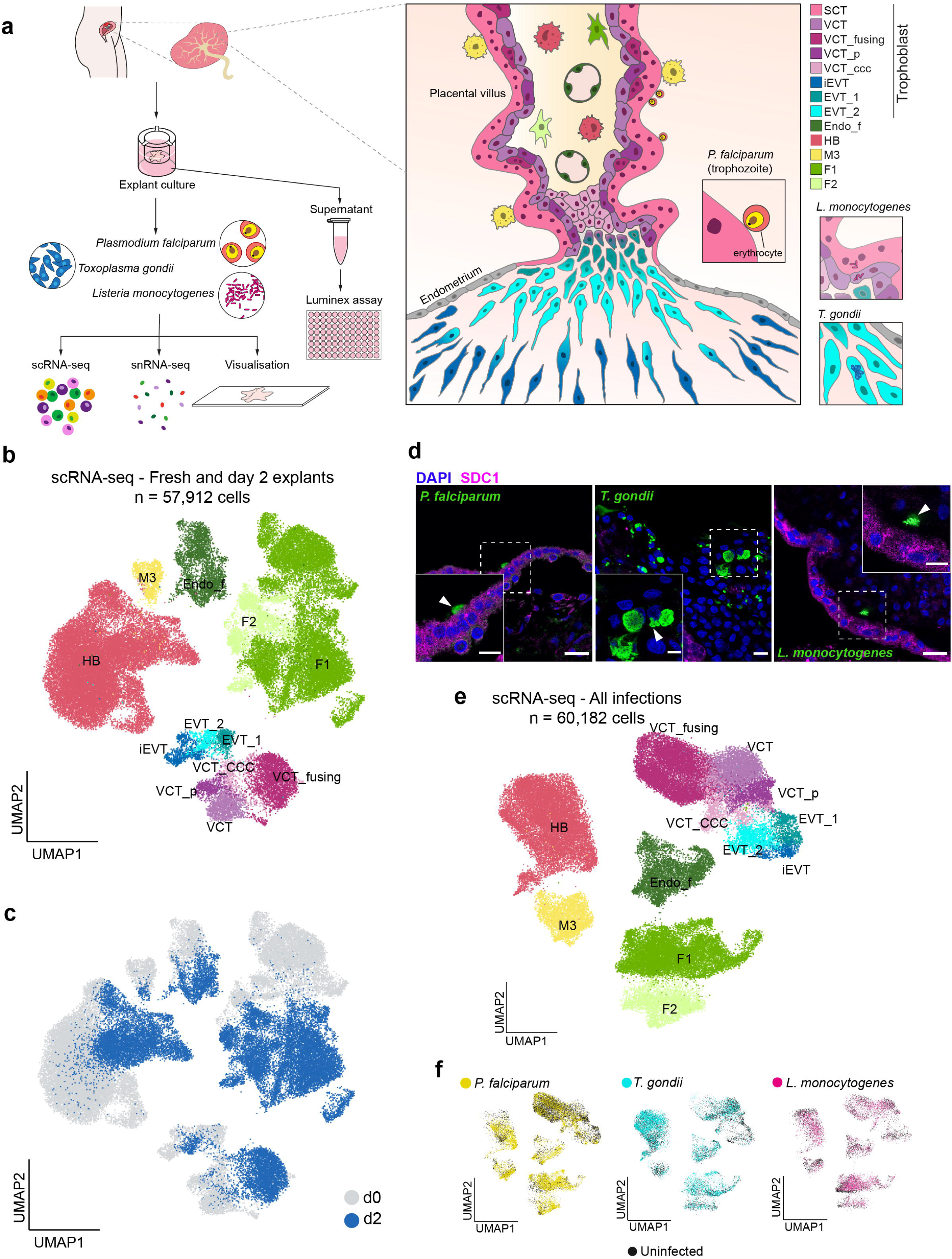
Placental explants in culture maintain cell lineages of the placenta and can be successfully infected with different pathogens. **a)** Left - Workflow of the study. Placental explants were grown in culture and infected with *P. falciparum, T. gondii* or *L. monocytogenes.* Samples were analysed using scRNA-seq, snRNA-seq and Luminex assay. Right - a schematic of a placental villus embedded in the endometrium showing all cell types and localisation of different pathogens. **b)** UMAP of cell lineages from fresh placental tissue and placental explants scRNA-seq (n=57,912) after 48 hours in culture. Colours correspond to the schematic in a). **c)** UMAP of time points from fresh placental tissue and placental explants scRNA-seq (n=57 912) after 48 hours in culture (gray - d0, blue - d2). **d)** Representative immunofluorescence staining of infected placental explants after 24h in culture, stained with SDC1 (magenta), a SCT marker, co-stained with DAPI (blue). Left - *P. falciparum* (green) attached to the outside of the SCT; middle - *T. gondii* tachyzoites (green) inside of the placental villus; right - *L. monocytogenes* (green) inside of the VCT. Arrowheads indicate individual parasites (*P. falciparum* and *T. gondii*) or a group of bacteria (*L. monocytogenes*). Scale bars: main panels - 20µm, insets - 10µm. **e)** UMAP of scRNA-seq (n=60,182) of uninfected and infected placental explants from matched donors after 24 hpi. Colours correspond to the schematic in a). **f)** UMAPs of of infected placental explants after 24 hpi showing the distribution of cells coming from experiments (controls uninfected and infected explants) with *P. falciparum* (yellow), *T. gondii* (cyan) and *L. monocytogenes* (magenta).

We first examined the heterogeneity of placental explants before and after cell culture by single-cell RNA sequencing (scRNA-seq) **(Supplementary Figure 1b-c)**. We collected 57,912 high quality cells of donor matched fresh placental cells and cells cultured for 48 hours **(Figure 1b, 1c)**. We integrated all conditions on the same manifold and annotated cell populations by transferring labels of our highly resolved map of the first-trimester placenta^35^ **(Supplementary Figure 3)**. After 2 days of culture in TOM, the placental tissue *ex vivo* culture system is capable of maintaining the major cell lineages: placenta-associated maternal macrophages (PAMM1), Hofbauer cells, trophoblasts, fibroblasts and endothelial cells **(Figure 1b, 1c)**. In addition, major intrinsic immune signatures, such as the expression of Toll-like receptors (TLRs) and interferon (IFN) receptors, are recapitulated in the macrophage and trophoblast populations after 48 hr of culture conditions **(Supplementary Figure 1e)**.

Next, we cultured three distinct regions of the placenta from two individuals and performed scRNA-seq analyses to ascertain the extent of heterogeneity of individual samples. Identity and proportions of all cell types are maintained across the different regions of the same placental sample, (Supplementary Figure 1d). This result means that the explant model is robust and its cell lineages and proportions are maintained when selecting distinct anatomical regions.

Altogether, placental explants are robust multilineage models able to recapitulate the heterogeneity of the first trimester placenta.

### ScRNA-seq and snRNA-seq of human placental explants infected with *P. falciparum, L. monocytogenes* and *T. gondii*

We next examined the impact of pathogen infection on placental explants, performing infections 12 hours after the start of the *ex vivo* culture. The explants were infected with extracellular (*P. falciparum*) and intracellular (*L. monocytogenes* and *T. gondii*) pathogens and cultured for 24 hours **(Supplementary Figure 1f, see Methods)**. We performed targeted adaptations of the media to promote viability of both host and pathogens, confirming the infectiousness and viability for all three pathogens in the defined culture medium **(see Methods)**. *P. falciparum* infections were performed in TOM supplemented with lipid-rich albumin (AlbuMAX) during the infection time-point^41^. For *L. monocytogenes* infections, we included gentamicin protection assay and defined TOM without antibiotic primocin^42^. For *T. gondii* infections, we used TOM without A83-01, an inhibitor of the TGF-β type I receptors ALK4/5/7.

For *P. falciparum* infection, we first enriched for parasites expressing the VAR2CSA *var* gene associated with placental malaria. Proteoglycan CSA is attached to several core proteins^43^ encoded by the following genes *CSPG2* (= *VCAN*), *CSPG4* and *CD44*, and its expression is conserved in the placental explants **(Supplementary Figure 4a)**. This enrichment is a key step as long term *in vitro* culture of *P. falciparum* elicits a heterogenous population of *var* genes expressed from the large gene family, with very low levels of *VAR2CSA* expression^44^. To facilitate parasite detection, we employed an eGFP-reporter line (PfNF54-eGFP)^45^ that was cycled through several rounds of panning on CSA to enrich for a *VAR2CSA*^high^ parasite population **(Supplementary Figure 4a)**. *VAR2CSA* expression was induced up to 50-fold after pan cycle 4, and maintained its expression after pan cycle 16, whereas expression of other *VAR* genes remained stable throughout the panning cycles **(Supplementary Figure 4b)**. This result supports the expansion of a clonal *VAR2CSA*^high^ specific population, which we termed as 2D9 PfNF54-eGFP. In addition, we also confirmed that the 2D9 *VAR2CSA*^high^ parasite population maintains endogenous expression of GFP at trophozoite stage **(Supplementary Figure 4c)**. We then utilised the synchronised ring stage 2D9 *VAR2CSA*^high^ parasites in all of our subsequent malaria infection assays.

Successful infection by each pathogen was confirmed by immunostaining **(Figure 1d)**. ScRNA-seq of all processed control and infected explants yields 60,182 cells after QC filtering **(Figure 1e, Supplementary Figure 3)**. Despite each pathogen having different mechanism of infection, syncytium-restricted for *P. falciparum*^16^ versus other trophoblast cell types for *L. monocytogenes*^12^ and *T. gondii*^14^, cells coming from explants infected with different pathogens are equally represented in all cellular clusters **(Figure 1f)**, indicating a lack of preferential cell survival or a technical artefact, such as differential cell survival after processing.

*P. falciparum* directly infects the external SCT layer and the primary response to malaria infection likely arises from the multinucleated syncytium, which warranted an in-depth investigation. To quantify the effects of infection on the multinucleated syncytial fraction of the trophoblast (SCT), which is lost by scRNA-seq, we performed single nuclei RNA sequencing protocol (snRNA-seq)**(see Methods)**. As a comparison with an intracellular pathogen infection, we also profiled explants infected with *L. monocytogenes*. After QC processing **(Supplementary Figure 3)**, we detect 18,496 nuclei, distributed over 8 trophoblast subpopulations^35^, including the SCT **(Figure 2a)**. As in our scRNA-seq data, nuclei coming from expants infected with different pathogens were represented in all cell clusters detected **(Figure 2b).**

**Figure 2.**
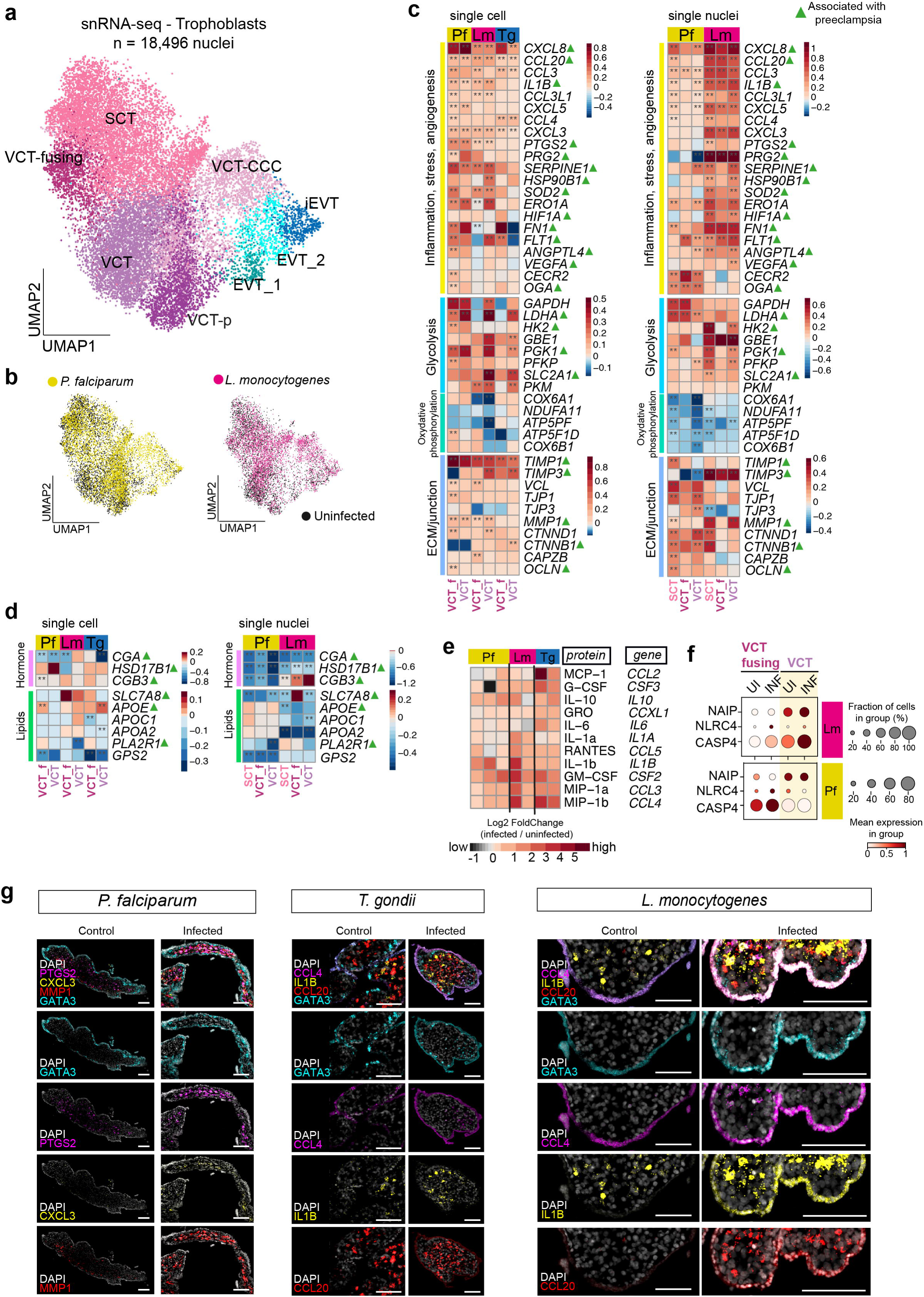
Trophoblast cells show signatures of robust inflammatory response upon infection. **a)** UMAP of trophoblast cell lineages from snRNA-seq (n=18,496) of uninfected and infected placental explants from matched donors after 24 hpi. Colours correspond to the schematic in Figure 1a. **b)** UMAPs of cells coming from infection experiments with *P. falciparum* (yellow) and *L. monocytogenes* (magenta) snRNA-seq. **c)** Heatmap showing log2fold changes of DEGs in different trophoblast cell lineages between uninfected and infected samples. Asterisk ** significance at adjusted *p* <0.05, Wilcoxon Rank Sum test. Genes have been divided into functional groups, based on their role in the placenta. Green triangles - genes associated with preeclampsia in the literature. **d)** Heatmap showing log2fold changes of DEGs involved in hormone and lipid metabolism in different trophoblast cell lineages between uninfected and infected samples (see Methods). Green triangles - genes associated with preeclampsia in the literature. **e)** Heatmap of log2fold changes of protein levels, detected through Luminex assay of supernatant form explant cultures after 24 hpi. Columns indicate biological replicates. **f)** Dot plots showing variance-scaled, log-transformed expression of inflammasome-related genes in scRNA-seq dataset from *P. falciparum*- and *L. monocytogenes*-infected explants. **g)** High-resolution large-area imaging of a representative placental explant section stained with *GATA3, PTGS2, MMP1* and *CXCL3* for *P. falciparum*-infected explant versus matched uninfected control and *GATA3, CCL3, IL1B* and *CCL20* for *T. gondii*- and *L. monocytogenes*-infected explant. Scale bars; 100 µm.

Thus, we have generated two manifolds in total: one manifold containing scRNA-seq data for *P. falciparum*, *L. monocytogenes* and *T. gondii* and another manifold containing snRNA-seq data for *P. falciparum* and *L. monocytogenes*.

### Trophoblast cells are capable of mounting an inflammatory response upon infection

We first studied the primary response of the trophoblast. Differential expression gene (DEG) and gene ontology (GO) enrichment analysis between infected and uninfected placental *ex vivo* tissue for individual pathogens reveals conserved signatures of response to infection in the main trophoblast populations in the placenta (SCT, VCT_fusing and VCT) **(Figure 2c, Supplementary Figures 5, 6)**. EVT subsets, which are located in the maternal endometrium *in vivo*, show considerably lower DEG **(Supplementary Figure 2e)**. Both upregulated and downregulated genes from each placental trophoblast subtypes (SCT, VCT_fusing and VCT) exhibits similar GO programmes between the scRNA-seq and snRNA-seq datasets in all infections **(Supplementary Figures 7, 8)**. This result indicated trophoblast are able to engage a common primary response to the three pathogens. Notably, *L. monocytogenes*-infected explants presented a higher number of DEGs, suggesting a stronger innate response to this bacterial pathogen.

Trophoblast cells are able to trigger an intrinsic inflammatory programme in response to infection^46, 47^. At 24 hours post infection (hpi), the trophoblast subtypes from infected placental explants upregulate inflammatory cytokine and chemokines genes (*IL1B*, *CXCL3*, *CXCL8*, *CCL3*, *CCL4* and *CCL20*) as well as the inflammatory gene cyclooxygenase *COX-2* (*PTGS2*)^48^ **(Figure 2c)**. We confirmed the strong upregulation of inflammatory cytokines and chemokines upon pathogen infection at the protein level by multianalyte profiling of the whole explant culture supernatant **(Figure 2e)**. Examples include an upregulation of IL1-b, MIP1-a and MIP1-b secreted protein levels, which correspond to the transcriptional levels seen in the scRNA-seq datasets **(Figure 2e)**. In agreement with the secretion of inflammasome-associated cytokine IL1-b upon infection, some inflammasome-associated genes including *NAIP, NLRC4 and CASP4*, are also found upregulated in our dataset **(Figure 2f)**^47, 49^.

In both single cell and single nuclei trophoblast datasets there is an increase in the expression of genes involved in glycolysis (for example *GAPDH, LDHA, GBE1, SLC2A1*), and a decrease in oxidative phosphorylation-related genes (*COX6A1, NDUFA11, ATP5PF*) **(Figure 2c)**. These synchronised changes might reflect a metabolic adaptation to increased demand for energy for cells undergoing an inflammatory response^50^. Specifically, increases in expression of *SLC2A1* (=*GLUT1*) and *GAPDH* are indicative of active glucose uptake and utilisation. In addition, our single nuclei data shows an increase in angiogenesis-related genes (*ANGPTL4, VEGFA, FLT1*) **(Figure 2c)**, which is another hallmark of a robust inflammatory response and is regulated by HIF1A^51^, which is also increased in the single nuclei dataset.

Our infection datasets show modest but significant increases in the expression of some adherent (*CTNND1, CTNNB1*) and tight (*TJP1, TJP3*) junction related genes in the trophoblast subsets **(Figure 2c)**. Increased expression of a matrix metalloproteinase *MMP1* underpins the remodelling of extracellular matrix (ECM), and has not previously been associated with infection by *L. monocytogenes*. Elevated expression of matrix metalloproteinase inhibitors - *TIMP*s (*TIMP1* and *TIMP3*) **(Figure 2c)**, especially in the *L. monocytogenes* infection experiments, further indicates the disbalance in MMP and TIMP ratios and is indicative of a robust inflammatory response in the trophoblast cells^52^. This is in concert with the dysregulation of ECM and cellular junctions^53^ in other inflammatory contexts.

To validate the expression of some inflammatory genes (*CXCL3, IL1B, CCL4, CCL20, PTGS2* and *MMP1*) in a spatial context and expression differences between pathogens, we performed multiplexed single-molecule fluorescent *in situ* hybridization (smFISH) in uninfected (control) versus infected placental explants at 24 hpi. *P. falciparum*-infected explant shows higher expression of *PTGS2, CXCL3* and *MMP1* within the villous core compared to uninfected control **(Figure 2g)**. *T. gondii*-infected explant shows higher expression of *CCL4* and *CCL20* particularly in the *GATA3+* trophoblasts epithelia **(Figure 2g)** in concordance with changes observed in our scRNA-seq and snRNA-seq datasets **(Figure 2c)**. *IL1B* expression remains localised at the villous core compartment in *T. gondii*-infected explant compared to uninfected control **(Figure 2g)**, likely expressed by the Hofbauer cells **(Figure 3b)**. In contrast, *L. monocytogenes*-infected explant shows significantly higher expression of *IL1B* and *CCL20* in the *GATA3+* trophoblast epithelia as well as in the inner villous core **(Figure 2g)**, further supporting that cytokine IL-1B may be involved in *L. monocytogenes* defence pathways^47^.

**Figure 3.**
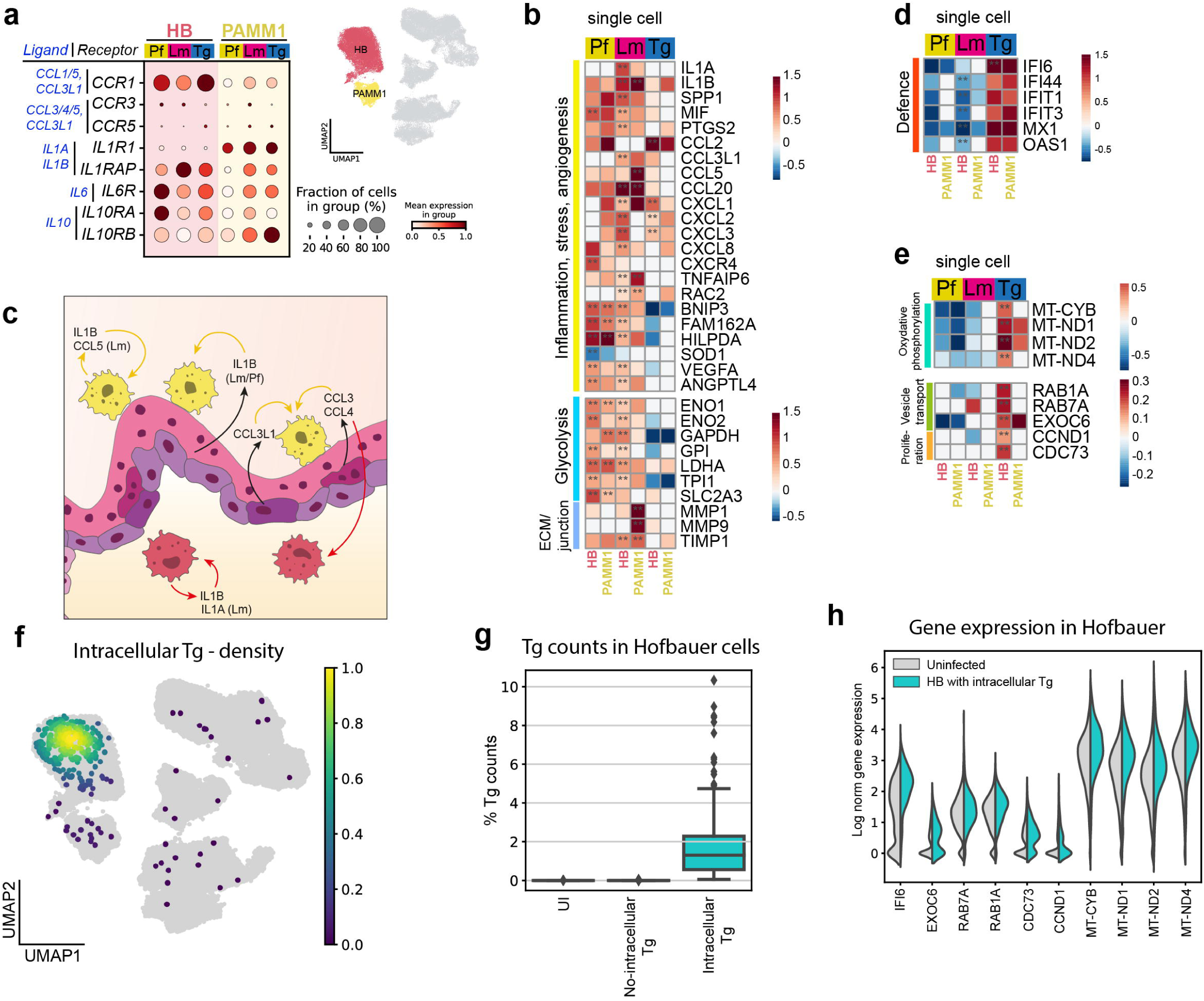
Maternal and fetal macrophages display pathogen specific immune programs with a generalised inflammatory response. **a)** Left - dotplot showing the expression of cytokine-receptor genes in Hofbauer (HB) and PAMM1 Macrophages. In blue the ligands (cytokines) expressed by trophoblast. Right - UMAP embedding highlighting the immune compartment. **b)** Heatmap showing log2fold changes of DEG in macrophages between uninfected and infected samples. Functional groups are based on their role in placenta. Asterisk ** significance at adjusted *p* <0.05, Wilcoxon Rank Sum test. **c)** Summary of trophoblast-macrophage and macrophage-macrophage communication mediated by the expression and sensing of cytokines. **d)** Heatmap showing log2fold changes between uninfected and infected samples of genes involved in defence responses. **e)** Heatmap showing log2fold changes between uninfected and infected samples of DEGs mainly affected by *T. gondii* infections. **f)** UMAP plot showing the density of cells with intracellular-*T. gondii* (*Intracellular-Tg*) in the embedding. **g)** Boxplot representing the percentage of *T. gondii* counts in Hofbauer cells. Uninfected samples (*UI*); Cells from Infected samples without detectable intracellular-*T. gondii* (*No-intracellular-Tg*); Cells with detectable intracellular-*T. gondii* (*Intracellular-Tg*). **h)** Violin plots showing the log-normalised expression of genes specifically upregulated after *T. gondii* infection.

Altogether, pathogen infection induces a primary inflammatory response in trophoblast, highlighting trophoblasts can mount a defence strategy for specific recruitment and activation of leukocytes to the trophoblast barrier.

### Trophoblast infection compromises placental function and has overlapping transcriptomic signatures with preeclampsia

Genes related to hormone pathways are prominently downregulated in our snRNA-seq dataset. The significant downregulation of *CGA,* and a similar trend, albeit to a lower degree, of downregulation of its partner *CGB3*, indicate a reduction in human chorionic gonadotropin (hCG) **(Figure 2d)**. hCG is one of the main players in the maintenance of pregnancy and the promotion of local immune tolerance. It is secreted by SCT^54^, which fits with the observed pattern of the greatest reduction observed in infection with the extracellular pathogen - *P. falciparum*. In addition, *HSD17B1*, which regulates the final step in estradiol biogenesis^55^ is downregulated by both *L. monocytogenes* and *P. falciparum* as shown by snRNA-seq **(Figure 2d)**. This may cause a reduced steroidogenic capacity of the placenta. Another group of downregulated genes are genes involved in transport and lipid metabolism. A reduction in the levels of a crucial amino acid transporter *SLC7A8* (=*LAT2*) combined with a downregulation of apolipoproteins (*APOE, APOC1, APOA2*), which are involved in the uptake of trophoblast of lipids^55^ point to an insufficiency in the transport function of the infected trophoblast.

Manual curation of DEGs in trophoblast subpopulations shows a high proportion of genes directly involved in preeclampsia **(Figure 2c - green triangles)**. Indeed, many of the processes identified in our infection model are disrupted in preeclampsia - inflammatory response (*ILB1, CCL20*)^56^, metabolism (*SLC2A1*)^57^, steroid production and hormone metabolism (*HSD17B1, CGA*)^58, 59^ and lipid metabolism and transport (*APOE*)^60^. These findings suggest that placental trophoblasts can mount robust inflammatory and barrier defence strategies at early infection stages. In parallel, placental trophoblasts undergo metabolic and functional adaptation in response to stress-induced inflammation, as evidenced by upregulation of glycolysis-related genes, potentially impacting its primary role as a nutrient-supplying and endocrine organ during early infection. Disruption of similar processes is observed in preeclampsia.

### Maternal and fetal macrophages elicit general and pathogen-specific immune responses to control infection

Macrophages dominate in the placental environment while adaptive immune cells are absent. There are two major placental macrophages: (i) PAMM1, maternal macrophages attached to the SCT that can aid in infection clearance and syncytium repair^34, 61^, and (ii) Hofbauer cells, fetal primordial macrophages that are located in the inner space of the placental villus. Cell-cell communication analysis using our tool CellPhoneDB^62^ shows that Hofbauer and PAMM1 macrophages potentially communicate with trophoblasts and between themselves via cytokine expression. Upon infection, Hofbauer and PAMM1 macrophages express receptors for inflammatory cytokines and chemokines **(Figure 3a)** which are produced by trophoblasts at 24 hpi **(Figure 2c, 2e)**. Those include *CCR1* (receptor of CCL1, CCL5, and CCL3L1), *CCR3* and *CCR5* (receptors of CCL3, CCL4, CCL5, and CCL3L1), *IL6R* (receptor of *IL-6*), *IL10RA* and *IL10RB* (receptors of *IL-10*) and *IL1RA* and *IL1RAP* (receptors of *IL1A* and *IL1B*) **(Figure 3a)**. In parallel, all three infections elicit an inflammatory response in macrophages that lead to the expression of cytokines and chemokines whose receptors are also expressed by the macrophages **(Figure 3b)**. This suggests trophoblast-macrophage and macrophage-macrophage signalling is happening. The potential interactions of this cellular communication are summarised in **Figure 3c**.

We identify metabolic adaptations in macrophages as a response to infection which may relate to the upregulation of an inflammatory response. Hofbauer cells and PAMM1 increase their glycolysis rates to satisfy the increased demand for energy during *L. monocytogenes* and *P. falciparum* infections, while there are no statistically significant changes identified in *T. gondii* infected samples. The increased expression of *SLC2A3* (=GLUT3), *GPI* (Glucose-6-Phosphate Isomerase) and *GAPDH* suggests an active glucose intake and metabolization in *L. monocytogenes* and *P. falciparum* infected samples. Hofbauer cells and PAMM1 also upregulate angiogenic genes (*ANGPTL4*, and *VEGFA*) in *L. monocytogenes* and *P. falciparum* infections **(Figure 3b)**. Other hallmarks of inflammation, including an elevated expression of *MMP1* and *MMP9*, and of the matrix metalloproteinase inhibitor *TIMP1* in PAMM1 upon Lm infection are also present in the macrophages **(Figure 3b, 2c)**.

*T. gondii* induces a distinctive transcriptomic profile in PAMM1 and Hofbauer cells **(Figure 3d, 3e)**. Hofbauer cells and PAMM1 significantly upregulate IFN inducible gene *IFI6* **(Figure 3d)**, and show an increased expression trend of other IFN-α/β genes (*IFIT1, IFIT1, IFI44*), suggesting that IFN-I responses are a key mechanism used by both macrophage populations to control *T. gondii* infection in the human placenta. In addition, Hofbauer cells increase mitochondrial activity, especially the subunits of the NADH dehydrogenase. The increase in NADH can favour the production of superoxide (O_2_^-^), which may have antimicrobial activity^63, 64^. Moreover, Hofbauer cells in *T. gondii*-infected samples show an increased expression of vesicular transport (*RAB1A, RAB7A* and *EXOC6*) and proliferation genes (*CCND1* and *CDC73*) **(Figure 3e)**. *T. gondii* parasite reads are mainly detected in Hofbauer cells, indicating the presence of intracellular *T. gondii* **(Figure 3f, 3g)**. The expression of the upregulated host genes during *T. gondii*-infection are visibly higher in Hofbauer cells with intracellular *T. gondii* compared to controls **(Figure 3h)**.

Altogether, we show PAMM1 and Hofbauer cells are key elements of the primary placental response to pathogens. We also show pathogen-specific effects in Hofbauer cells in response to *T. gondii* infection.

### Placental infection alters erythrocyte remodelling, nutrient acquisition and metabolism programs in the human malaria parasites

We have defined multiple adaptations of the placental microenvironment in response to pathogens. Next, we explored how the pathogen also adapts to changes in its microenvironment by looking at *P. falciparum*, which is known to induce phenotypic and physiological adaptations in response to different host niches^65, 66^. To profile local adaptation events of the human malaria parasite upon placental-tissue encounter, we performed scRNA-seq on *Plasmodium* infected erythrocytes - 2D9 *VAR2CSA*^high^ isolated from placental explants **(Figure 4a)** and compared them to their non-infected counterparts on the same donors profiled for host responses **(see Methods)**. In total, three parasite fractions were generated from each placental infection assay: (i) ‘pf_b’ (binder) comprising cytoadherent *VAR2CSA*^high^ population bound to the placenta, (ii) ‘pf_nb’ (unbound fraction) comprising of a washed fraction (i.e. parasite exposed to the placenta culture but not tightly bound to it at the moment the tissue was collected) and (iii) ‘pf_iv’ (*in vitro)* comprising of the same parasite batch and stage that were used for infection, cultured in parallel in equivalent medium but not exposed to human placenta **(Figure 4a)**. We then used flow cytometry to measure parasite viability and cell counts **(Supplementary Figure 9a)**. The washed fraction ‘pf_nb’ has a similar cell count to the ‘pf_iv’ sample, but the placental isolate ‘pf_b’ has a higher cell count, supporting an enriched-binding phenotype **(Supplementary Figures 9a-b)**. To identify parasite gene signatures associated with the binding phenotype, we subjected the parasite fractions to scRNA-seq.

**Figure 4.**
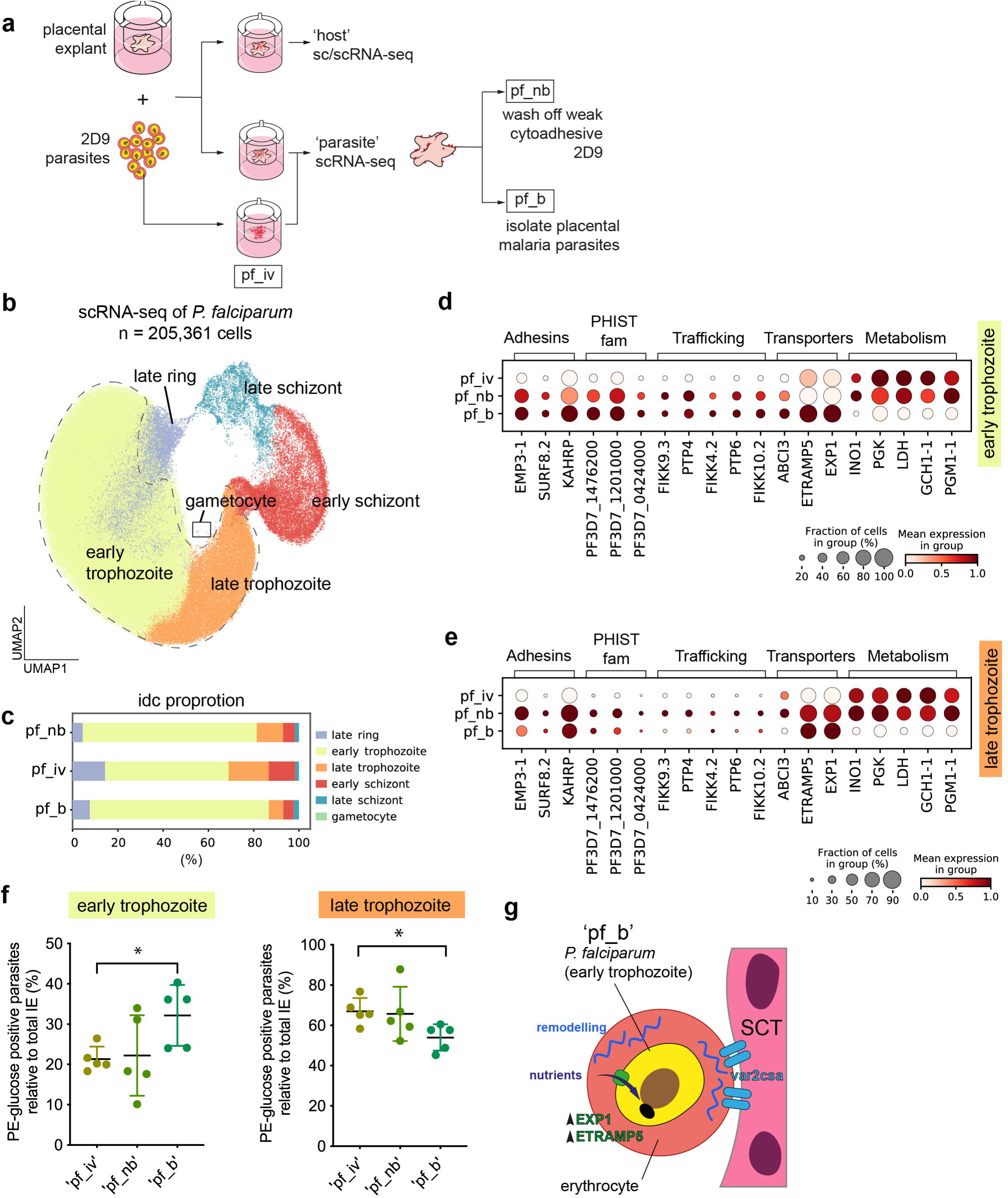
Placental infection alters erythrocyte remodelling, nutrient acquisition and metabolism programs in the human malaria parasites. **a)** Schematic of the workflow for *P. falciparum* culture and explant infections. **b)** UMAP of *P. falciparum* life cycle scRNA-seq (n=205,361). **c)** Bar plots showing proportion of *P. falciparum* life cycle for different conditions (pf_nb, pf_iv, pf_b). **d)** Dot plots of significant DEGs (*p* adj < 0.05) between ‘pf_b’ and ‘pf_iv’ at early trophozoite stage. **e)** Dot plots of the same list of genes from (d) but at late trophozoite stage. Dot plots are variance-scaled and log-transformed. **f)** Percentage of PE-glucose positive parasites relative to total number of infected erythrocytes at early trophozoite (left panel) and late trophozoite (right panel) stage. Asterisk * significance at *p* <0.05 using non-parametric Mann-Whitney test, n=5. **g)** Schematic diagram to illustrate ‘pf_b’ adaptive change upon cytoadherence to human placental tissue.

After quality control steps **(Supplementary Figure 10a)**, we generated a total of 205,631 single-cell transcriptomes of human malaria parasites from three independent placental explant infections **(Figure 4b, Supplementary Figure 10b)**. We used intraerythrocytic developmental cycle (IDC) marker expression on Leiden clusters to map the intraerythrocytic developmental stages of the parasite population **(Supplementary Figure 10c)**. As expected, the majority of the 2D9 *VAR2CSA*^high^ parasites captured are assigned to early and late trophozoite stages with a small proportion of schizont and ring stages **(Figure 4b, 4c)**. *VAR2CSA* gene expression peaks at early-ring to mid-trophozoite stages **(Supplementary Figure 10d)**. The trophozoite is the primary cytoadherence stage at the placenta, as this stage corresponds to the peak of VAR2CSA protein expression^67^. Thus, we subsetted the early- and late-trophozoite clusters for DEG analysis between ‘pf_b’ and the ‘pf_iv’ control **(Supplementary Figure 10e-f)**.

At the early trophozoite stage, parasites isolated from placental explants (‘pf_b’) upregulate genes associated with erythrocyte cytoskeleton remodelling (*KAHRP, EMP3, SURF8.2*), the *PHIST* family members (*PF3D7_1476200, PF3D7_1201000*) and trafficking of PfEMP1 surface adhesins (*PTP4, PTP6, FIKK10.2, FIKK9.3, FIKK4.2*) **(Figure 4d)**. These changes may further facilitate cytoadherence of the parasites to the host placental tissue. To a lesser extent, the washed fraction ‘pf_nb’ recapitulates some of the gene expression features observed in ‘pf_b’. This is likely due to the fact that ‘pf_nb’ includes both parasites that are weakly bound to trophoblasts as well as non-binders. By contrast, parasite glycolytic metabolism genes (*PGK, LDH, PGM1*) are significantly downregulated in ‘pf_b’ **(Figure 4d)**. Upon transition to the late trophozoite stage, differences for gene categories involved in adhesin, nutrient transport and metabolism in ‘pf_b’ are consistent **(Figure 4e)** yet the changes in upregulated genes between ‘pf_b’ and ‘pf_iv’ control are more modest **(Supplementary Figure 10f)**.

The nutrient transporter genes (*EXP1, ETRAMP5, ABCI3*) are significantly upregulated in ‘pf_b’ in the trophozoite stage **(Figure 4d, 4e)**. EXP1 is known to mediate nutrients uptake such as glucose into *P. falciparum* trophozoites and we validated this using a fluorescence-labelled glucose uptake assay in a *P. falciparum* EXP1 knockdown line (Pf3D7-EXP1wt^low^-Ty1 (*sfa32*)) **(Supplementary Figure 11 b-e)**^68^. To demonstrate that the interaction with placental tissue also alters nutrient uptake by the parasite, we performed the fluorescence-labelled glucose uptake assay on ‘pf_b’ compared with the ‘pf_iv’ and ‘pf_nb’ parasite fractions. We calculated the percentage of PE-glucose positive parasites out of the total infected erythrocyte population for each parasite sample. ‘pf_b’ shows an increase in glucose uptake compared to ‘pf_iv’ control in early but not late trophozoite population **(Figure 4f)**. This increase in glucose uptake is likely to be induced by the upregulation of nutrient transporters in the early trophozoite ‘pf_b’ parasites. These findings suggest a new way for the human malaria parasite to adapt to the placental microenvironment.

In sum, placental infection induced transcriptional changes in the malaria parasites with key changes in genes involved in erythrocyte remodelling, protein trafficking, nutrient acquisition and glycolytic metabolism. It further suggests that malaria parasites can adapt in response to the host tissue type.

## Discussion

The placenta is the extra-embryonic interface that protects and nourishes the embryo during pregnancy. It is established early during development and functions as a barrier tissue that protects the embryo lacking adaptive immune responses ^22, 69^. Despite the relevance of the placenta in the nourishment and protection of the embryo, the impact of infections on the function and innate immune response of first trimester placenta are unclear. Here we characterise the primary response of the human placenta to infections during early human development by using an improved placental explant culture technique. Systematic characterisation of the multiple cell lineages using single cell and single nuclei profiling demonstrate that first-trimester placental explants have similar cellular composition to the *in vivo* placenta. We then utilise this culture system to understand the host-pathogen response when challenged with relevant infectious agents to model the features of placental malaria, toxoplasmosis and listeriosis. We demonstrate that the placental explant culture can be successfully infected with *P. falciparum*, *T. gondii* and *L. monocytogenes*, revealing coordinated host responses by trophoblast and fetal macrophages as well as parasite adaptations to the placental environment.

The trophoblast exerts a systemic inflammatory response upon infection with all three classes of pathogens. Increased expression of chemokines such as *CCL3* (MIP1a) and *CCL4* (MIP1b) at the transcriptional and protein level may activate the rapid mobilisation of maternal leukocytes to the site of infection and inflammation, directly from maternal blood. These responses to infection reflect a host innate immune defence strategy to minimise impending tissue injury and spread of infection^70^. Despite serving as a protective strategy, this inflammation response disrupts the core functions of the trophoblast, particularly hormone and lipid metabolism, which could have profound impact on the transport of nutrients to the embryo and on fetal growth and organ development^71–73^. Inflammatory and stress conditions can induce a switch in metabolic demands^74^. We observe that trophoblast and immune cells infected with *P. falciparum* and *L. monocytogenes* preferentially upregulate glycolysis over oxidative phosphorylation-associated genes, suggesting a transition to a metabolically active state necessary for the production of biomolecules such as cytokines and chemokines.

Concurrent with inflammatory and chemotactic events, we discovered a dysregulation in the hormonal milieu of the infected placenta that could have significant impact on pregnancy outcomes and fetal development. Placental hormone hGC (composed of subunits coded by *CGA* and *CGB3*) imbalance is associated with intrauterine growth restriction^75, 76^. These correlations of the disruptions in individual core functions of the placenta with pregnancy complications suggest the redirection of placental resources into inflammation response during infection detracts from the direct embryo-supporting functions, diminishing the access of the embryo to nutrients and hormones.

Surprisingly, a lot of DEGs observed in our infection model overlap with the genes associated with preeclampsia^77^. Disruption of similar processes in both infection and during aetiology of preeclampsia might explain the observed overlap in gene expression. It could also provide insight into correlations of post-infection preeclampsia observed in some studies^78–81^.

In other systems, pathogens modulate the ECM and tight junction to enable easier entry into the cell, and the host remodels it in order to recruit immune cells and facilitate infiltration ^53^. Since the trophoblast does not permit infiltration of maternal immune cells, dysregulation of its ECM and cellular junctions might serve as a signal for recruitment of immune cells in response to infection. Moreover, the trophoblast layer likely plays a role in immune-regulation through cell-to-cell communication with nearby structural and immune cells^82^, such as PAMM1 and Hofbauer cells. For instance, the *IL1B* expression in trophoblasts during *L. monocytogenes* infection might prime PAMM1 macrophages, which express the cognate receptors ILR1 and IL1RAP. Thus, trophoblast structural and inflammatory dysregulation in the context of infection may have a direct impact on immune cell function in the placental microenvironment as well^83^.

We classified generalised immunomodulation features in the maternal PAMM1 and Hofbauer cells, likely contributing to the overall inflammatory state of the tissue. Hofbauer cells have been recently demonstrated to have defence roles, including phagocytosis and formation of acidic phagosomes^61^. Intriguingly, *T. gondii* parasite reads from scRNA-seq are detected predominantly in the Hofbauer cells, suggesting that Hofbauer cells may have phagocytic activity for intracellular *T. gondii. T. gondii*-infected Hofbauer cells displayed a positive regulation in mitochondrial encoded genes, further supporting a dysregulation in mitochondrial metabolic activity^84^. These changes might reflect a direct interaction between *T. gondii* and the host mitochondria as *T. gondii* can hijack the functions of the outer mitochondrial membrane (OMM) through direct contact^85^. The increase in vesicular transport and proliferative genes in *T. gondii*-infected Hofbauer cells suggests that *T. gondii* may co-opt these tissue resident macrophages for replication and transmission to neighbouring cells, similar to the mechanism proposed to happen during Zika virus infection^86^, where Hofbauer cells play a key role spreading the virus.

Understanding how the pathogen responds to changes in the host is a crucial aspect of host-pathogen biology. Our current experimental strategy has enabled us to sequence high quality *P. falciparum* reads and profile how human malaria parasites respond to the host placenta. *Plasmodium* are known to manifest adaptation features when exposed to challenging host microenvironments such as the bone marrow, spleen or liver^65, 66, 87^. Rapid adaptation of the malaria parasite is likely crucial for its survival, replication and transmission. We observe that *P. falciparum*, when exposed to the human placenta through its interaction with SCT, showed increased expression of adhesins and erythrocyte remodelling gene categories. These genes encode factors that favour parasite adhesion that would eventually promote cytoadherence to the host epithelial and endothelial lining^88, 89^. *P. falciparum*-infected erythrocytes are prone to systemic clearance by the spleen and hence, the parasite has evolved its cytoadherence strategy to escape immune clearance^88, 89^. We observe elements of this unique mechanistic property of malaria parasites for the first time for placental malaria. In the presence of placenta, we further propose that *P. falciparum* faces nutritional and metabolic challenges^90^, thus triggering its nutrient efflux and metabolic switch properties. Understanding the adaptive strategies employed by the parasite during placental malaria may potentiate the design of novel and safe antimalarial drugs specific for pregnant women in malaria endemic countries^91^.

Overall, we provide the first detailed insight into the primary responses of the human placental trophoblast and macrophage against acute infections during human early development. Our study also provides mechanistic insights into host-pathogen interaction dynamics at the human maternal-fetal interface.

## Methods

### Samples

All tissue samples were obtained with written informed consent from all participants in accordance with the guidelines in the Declaration of Helsinki 2000. Placental samples were obtained from elective terminations from The MRC and Wellcome-funded Human Developmental Biology Resource (HDBR, http://www.hdbr.org), with appropriate maternal written consent and approval from the Fulham Research Ethics Committee (REC reference 18/LO/0822). The HDBR is regulated by the UK Human Tissue Authority (HTA: www.hta.gov.uk) and operates in accordance with the relevant HTA Codes of Practice.

### Placental explant culture

Samples of placental tissue were collected from surgical terminations between 5 and 17 post-conceptual weeks (pcw). After collection, samples were kept on ice in Ham’s/F12 media, shipped to the institute and processed within eight hours of collection. Samples were washed in ice-cold Ham’s/F12. The whole sample was cleaned of blood clots and the villi identified. The area containing the most villi was cut using a scalpel to size of approximately 1 cm^2^. ThinCert™ 6-well plate inserts (Greiner bio-one, 657640) were pre-coated with 50% Matrigel® (Corning, CLS356255) in ice-cold Trophoblast Organoid Media (TOM) supplemented with 10%FBS (ref. Turco 2018.) (see below). After polymerisation, placental explants were transferred to ThinCerts, the 6 well plate was filled with warm TOM (Advanced DMEM/F12 (Gibco, 12634-010), 1X B27-Vitamin A (Life Technologies, 12587010), 1x N2 (Life Technologies, 17502048), 100 µg/mL Primocin (Invivogen, ant-pm-1), 1.25 mM N-Acetyl-L-cysteine (Sigma-Aldrich, A9165-5G), L-glutamine (Sigma-Aldrich, 25030-024), 50 ng/mL recombinant human EFG (Peprotech, AF-100-15), 1.5µM CHIR99021 (Tocris, 4423), 80 ng/mL recombinant human R-spondin-1 (R&D Systems, 4645-RS-01M/CF), 100 ng/mL recombinant human FGF-2 (Peprotech, 100-18B), 50 ng/mL recombinant human HGF (Peprotech, 100-39), 500nM A83-01 (Tocris, 2939), 2.5 µM prostaglandin E_2_ (Sigma-Aldrich, P0409), 2 µM Y-27632 (Millipore, 688000)) and placed in the incubator at 37°C (21% O_2_, 5% CO_2_). Explants were infected after ∼12 hr (hour) in culture and processed after 24 hr and 48 hr post infection. For intra sample variability detection, explants were processed after 24 hr in culture.

### Culture of L. monocytogenes

A single colony of *L. monocytogenes* EGD-cGFP strain^92^ was picked from streaked Brain Heart Infusion (BHI) agar plate and inoculated in 10 ml of BHI broth. BHI broth was cultured overnight for 15hr in slanting position at 37°C at 120 rpm. EGD-cGFP optical density at 600nm (OD_600nm_) was measured using a spectrophotometer (Biowave CO8000) and re-diluted to OD_600nm_ 0.1. EGD-cGFP was then cultured in fresh BHI for another 2-3 hr until mid-exponential growth phase prior to placental explant infection.

### Culture of *T. gondii* and tachyzoite isolation

Human foreskin fibroblasts (HFFs) (ATCC, ATCC-CRL-2429) were maintained in D10 media (DMEM, high glucose (Gibco, 41965062), 10% FBS, 1% L-glutamine (ThermoFisher Scientific, 10500064), 0.5% penicillin/streptomycin and 0.1% Amphotericin B (Thermo, 15290026)) and passaged by using Tripsin-EDTA 0.25% (Gibco, 25200072). Confluent HFFs were infected with the RH+HX strain of *T. gondii* in ED1 media (DMEM, high glucose, 1% FBS, 0.1% L-glutamine, 0.5% Pen/Strep, and 0.1% Amphotericin B). The *T. gondii* culture was propagated 2-3 times a week by infecting a new batch of HFFs.

To isolate and synchronise tachyzoites, the media of infected HFF cultures of ∼40 hpi (with visible parasitic vacuoles) was removed and cells were washed 2 times with 1x PBS to remove extracellular parasites. D10 media was added to the HFFs, a cell scraper was used to detach the cells. The detached cells were transferred into a syringe with a 27 gauge needle and poured into a 15ml falcon tube, the pressure created while passing the cells through the needle bursts the cells and releases the intracellular parasites. The parasite/host suspension was filtered using a 20µm filter and then centrifuged at 100g for 5 min (minutes) to pellet host cell debris. The supernatant was centrifuged at 900g for 15 min to pellet the isolated tachyzoites.

### Culture of human malaria parasite *Plasmodium falciparum* and *in vitro* selection procedures

*P. falciparum* NF54-eGFP^45^, 2D9 NF54-eGFP and *P. falciparum* 3D7-EXP1wt^low^-Ty1 (*sfa32*) in condΔEXP1^68^ were maintained in 3% hematocrit in human O+ blood in RPMI 1640 supplemented with 5% Albumax-II and 50µg/ml hypoxanthine (Gibco), 25mM HEPES (Sigma) and 1x L-Glutamax (Gibco) in mix gas containing 5% O_2_, 5% CO_2_ and 90% N_2_. Human O+ blood was provided by the NHS Blood and Transplant, Cambridge, UK. NF54-eGFP were synchronised at high parasitemia levels by 70% Percoll (17-0891-01, VWR) gradient and 5% sorbitol lysis as reported previously^93^. Selection for *VAR2CSA*^high^ was done by repeated panning of trophozoite-infected erythrocytes (IE) on 150mm petri-dishes pre-coated with chondroitin sulphate A (CSA) sodium salt from bovine trachea (Merck) ^94^. The resulting parasite line after repeated panning was designated as 2D9. Ring stage parasites of 3D7-EXP1wt^low^-Ty1 (*sfa32*) in condΔEXP1 were synchronized twice with 5% sorbitol lysis within a 5 hour developmental stage window. Parasite cultures were split into two flasks and 250 nM rapalog (AP21967, Clontech) was added to one of the two flasks to induce excision of endogenous EXP1, resulting in EXP1^low^ parasites with an episomal copy of EXP1 driven by the *sfa32* promoter. The flask without rapalog treatment served as a wild-type control culture. The medium containing rapalog was changed daily and the parasites further cultured in the presence of rapalog for 18-24 hr where trophozoite stages were used for the glucose uptake assay.

### *var2csa* expression analysis by quantitative Real-Time PCR

Total RNA from synchronised ring stage 2D9 PfNF54-eGFP and unpanned bulk PfNF54-eGFP cultures were isolated using Trizol-chloroform (Invitrogen) as previously reported^95^. The total RNA was further purified using on-column PureLink® DNase Treatment (12185010, Invitrogen) according to the manufacturer’s instructions. 1µg of purified RNA was subjected to cDNA synthesis with SuperScript III First-Strand synthesis system (18080051, Invitrogen) and primed with Oligo(dT)20 primers (18418020, Invitrogen), random hexamers (48190011, Invitrogen) and dNTP mix (18427013, Invitrogen) in a 20µl reaction volume according to the manufacturer’s recommendation. A cDNA synthesis reaction without reverse transcriptase enzyme was included as control. 10-fold diluted cDNA was used for quantitative Real-time PCR with Power SYBR Green PCR Master Mix (4367659, Invitrogen) master mix with 500nM forward and reverse primer pairs (Supplementary table 1) in a final volume of 20µl. Reactions were run on the StepOnePlus Real-Time PCR Systems (Thermo Fisher Scientific) at default parameters; 95℃ for 10 min, then subjected to 40 cycles of 95℃ for 15 s and 60C for 1 min, with a subsequent melting step. Transcriptional level was normalised against the housekeeping gene *PF3D7_1218600* (arginine-tRNA ligase). Relative gene expression was calculated by 2^-ΔΔCt^ method, with a different 2D9 PfNF54-eGFP panned cycles compared to unpanned bulk PfNF54-eGFP culture.

### Infection of human placental tissue explants

All placental explants were washed three times with warm 1xPBS prior to infection with pathogens. Infection was done in duplicate wells per donor tissue, where one well was processed for single-cell isolation and the other was cryopreserved for single-nuclei isolation After infection, culture supernatant was collected and snap frozen for each of the pathogen infection and time-points.

#### L. monocytogenes

Mid-exponential phase *L. monocytogenes* culture was washed twice with TOM/10%FBS without primocin. Placental explants were infected with 1.5 x 10^8^ CFU of *L. monocytogenes*^42^ for 5 hr in TOM without primocin antibiotic. Then the explants were washed three times with 1x PBS before adding in TOM with 50 µg/ml of gentamicin (15750060, Thermo Fisher) for 24 hr. Placental tissue was washed again three times with 1x PBS and subjected to either single-cell processing or cryopreservation.

#### T. gondii

All infections with *T.gondii* were performed using TOM/10%FBS *without* A83-01 (TOM -A83), as we found that this compound impedes the infection process by restricting the parasite replication (data not shown). Placental explants were infected with 1 x 10^6^ isolated tachyzoites for 5 hr in TOM -A83. Then the explants were washed three times with 1x PBS, TOM -A83 was added to the washed explants for 24 hr. Placental tissue was washed again three times with 1x PBS and subjected to either single-cell processing or cryopreservation.

#### P. falciparum

Placental explants were infected with 300,000 parasites per µl of erythrocytes according to the World Health Organization (WHO) definition of hyperparasitemia (above 250,000 parasites per µl of erythrocytes)^96^. The explants were washed three times with 1x PBS before adding in parasites resuspend in TOM with 5% KnockOut Serum Replacement containing lipid-rich albumin (AlbuMAX) (Gibco, 10828028) for 24 hr. Parasites and explant co-culture, including the ‘pf_iv’ controls were maintained at 37°C with mammalian culture gas mix 21% O_2_, 5% CO_2_. Parasites viability and growth were validated in these gas mixtures within the 24 hr culture window. After infection, placental tissue was washed again three times with 1x PBS and subjected to either single-cell processing or cryopreservation.

### Processing of placental tissue explants for scRNA-seq

Explants were treated with Corning® Cell recovery solution (Corning, CLS354253) for 15 minutes on ice to remove Matrigel® remnants. The plate was shaken every 5 minutes to dislodge the explant from the membrane. The tissue was transferred to a 50mL Facon tube with ice-cold PBS and washed by inverting, then transferred to a Petri dish with fresh ice-cold PBS, and the villi were gently scraped using forceps and scalpel. Using a Pasteur pipette with a cut tip, all of the tissue was transferred to a Falcon tube with 40 mL of pre-warmed 0.2% trypsin-EDTA and sealed with parafilm. Digestion was performed at 37°C on a rotator with gentle rotation at 30 rpm for 7 minutes and stopped with 2mL of PBS. The tissue was filtered using a 100µm strainer and the strainer was washed with 5-10 mL of RT PBS (the tissue on the filter was kept for the second step of digestion). The filtrate was centrifuged at 200*g* for 5 minutes at RT. The supernatant was discarded and the pellet kept on ice (digest 1). The undigested tissue was retrieved from the filter by forceps and placed in a 50 mL Falcon tube containing 12.5mL of prewarmed 1mg/ml Collagenase V (Sigma-Aldrich, C9263) and 40µg/mL DNase I (Roche, 10104159001) in PBS. The tube was sealed with Parafilm and transferred to a tube rotator in the incubator. The tissue was digested for 30 minutes with slow rotation at 30 rpm. The cell suspension was filtered through a 100µm filter and the filter was washed with 5mL of RT PBS. The filter with any undigested tissue was discarded. Cells were pelleted at 200*g* for 5 minutes at RT (digest 2). Cells from digest 1 and digest 2 were pooled and washed with PBS with 20µg/mL of DNase I (DNase I buffer). Cell suspension was centrifuged at 200*g* for 5 minutes at RT, the pellet resuspended in 1mL of DNase I buffer and kept on ice. Cells were counted using a haemocytometer.

To remove erythrocytes and debris, a Pancoll gradient was used. The cell suspension was topped up to 10mL with DNase I and very slowly added on top of 5mL of Human Pancoll (PAN-Biotech, P04-60500) in a 15mL Falcon tube. The suspension was centrifuged at 600*g* for 20 minutes at RT, with using acceleration=0, break=1 settings on the centrifuge. Cells were collected from the interface using a Pasteur pipette. The cells were washed with DNase I buffer and centrifuged at 200*g* for 5 minutes at RT twice. Cells were passed through a 35-40µm filter and counted (bulk cells). 1 million of these bulk cells were collected for flow cytometry assessment (see Flow cytometry section).

To enrich the final cell suspension for the CD45+ population, 1 million of bulk cells were separated and kept on ice and the rest was submitted to the MACS-based enrichment following the CD45 MicroBeads protocol (product code). The cells were centrifuged at 200*g* for 5 minutes at RT and the pellet resuspended in DNase I buffer and counted (CD45 cells). Bulk cells and CD45 cells were mixed in 1:1 ratio in PBS/0.04% BSA prior to loading of the 10x Genomics Chromium inlet.

Three independent placental tissue regions per donor were excised and processed to single cell suspension. For hashtag antibody staining, 500,000 cells from bulk and CD45+ enriched fraction were individually re-suspended in Cell Staining Buffer (420201, Biolegend) and blocked with 5 µL Human TruStain FcX™ (422301, Biolegend) for 10 minutes at 4 °C. Each of the cell suspensions originating from different placental tissue regions were hashtagged with different TotalSeq-B™ antibodies for 30 minutes at 4 °C. 1 µg of the following TotalSeq-B™ antibodies were used; TotalSeq™-B0253 anti-human Hashtag 3 Antibody (394635, Biolegend), TotalSeq™-B0254 anti-human Hashtag 4 Antibody (394637, Biolegend) and TotalSeq™-B0255 anti-human Hashtag 5 Antibody (394639, Biolegend). Antibody-tagged cells were washed three times with 3 ml of Cell Staining Buffer using centrifugation speed of 500 x g for 5 minutes at 4°C. Bulk cells and CD45 cells were mixed in 1:1 ratio then filtered through a 40 uM cell strainer. Cells were counted and diluted to desired loading concentration to recover between ∼4000 cells. Hashtagged cells were pooled at equal cell number ratio (targeted for a total of ∼12000 cells) right before loading of the 10x Genomics Chromium inlet.

### Isolation of *P. falciparum* 2D9-IEs from the placental tissue explants for scRNA-seq

Infected placental tissue explant with 2D9 PfNF54eGFP were washed on a rotating platform for five times with 1xPBS to remove unbound parasites, which was designated as ‘pf_nb’. The washed placental tissue explants were then cut into smaller pieces, followed by another gentle washing with 50 mM EDTA (15575020, Invitrogen) on a rotating platform for 1 hr, as previously described^97^. The EDTA washed fraction, containing potentially cytoadherence population, was filtered through 40µM cell strainer to remove placental tissue pieces. The filtered fraction containing mostly bound parasites, designated as ‘pf_b’, was centrifuged at 1500 rpm and washed five times with 1xPBS. 2D9 PfNF54eGFP used for infection assay was also cultured in parallel in the same batch of TOM media, but without the exposure to placental tissue explants. This batch of culture is designated as ‘pf_iv’ which was also harvested and washed 5 times with 1xPBS. All three parasite fractions, ‘pf_b’, ‘pf_nb’ and ‘pf_iv’ were then resuspended in PBS/0.04% BSA at 3% hematocrit for flow cytometry assessment and loading of the 10x Genomics Chromium inlet.

### Processing of samples for snRNA-seq

Single-nuclei suspensions were isolated either from fresh frozen placental explants or OCT frozen tissue sections (8 x 50 μm cryosections) using a previously described protocol^98^. Isolated nuclei were stained with Trypan blue and counted with a haemocytometer. Nuclei suspension was then resuspended in PBS/1% BSA diluted to targeted final concentration to recover between ∼5,000-6000 nuclei per sample prior to loading prior to loading of the 10x Genomics Chromium inlet.

### 10x Genomics Chromium gene expression library preparation and sequencing

Cells and nuclei were loaded into the 10x Genomics Chromium inlet according to the manufacturer’s instructions. The Chromium NEXT GEM Single Cell 3′ Library v3.1 chemistry (10x Genomics) was used and libraries were prepared according to manufacturer’s instructions aiming to recover between 2,000 and 25,000 cells per reaction. For cell hashing workflow, Chromium NEXT GEM Single Cell 3′ Library v3.1 with Feature Barcoding technology for Cell Surface Protein was used according to the manufacturer’s protocol (10x Genomics). Sequencing was done at a minimum coverage of 20,000 read pairs per cell for gene expression libraries and 5000 read pairs per cell for cell surface protein libraries on an Illumina Novaseq S4 platform.

### Flow cytometry

#### Placental tissue

Bulk single cell suspension was filtered through a 40 uM cell strainer, then washed in the FACS buffer [2%FBS/2mM EDTA in PBS]. Cells were blocked with Human TruStain FcX™ (422301, Biolegend) for 10 minutes at 4 °C. Cells were stained with 1:40 FITC anti-human CD45 antibody (304006, Biolegend) and 1:400 LIVE/DEAD™ Fixable Near-IR Dead Cell Stain (L10119, Thermo Fisher) for 20 minutes at 4 °C. Cells were washed twice with FACS buffer by centrifugation at 5 minutes 200x g at 4 °C before a final resuspension in 300 µl of FACs buffer. Flow cytometry was performed using the CytoFLEX cytometer (Beckman Coulter). Flow cytometry datasets were analysed by FLOWJO Software v10.6.1 (Tree Star).

#### Malaria parasites

5µl of IE at 3% hematocrit with endogenous GFP expression were labelled with 95µl staining solution containing 200nM of Mitotracker Red FM (M22425, Invitrogen) and/or 100nM of SYBR Green (S7563, Thermo Fisher) for 20 minutes at 37°C. Mitotracker dye stock was dissolved in DMSO at a concentration of 1mM prior dilution to working concentration. Uninfected erythrocyte was included as background gating control. Flow cytometry was performed using the CytoFLEXcytometer (Beckman Coulter). Flow cytometry datasets were analysed by FLOWJO Software v10.6.1 (Tree Star).

### Tissue cryopreservation

Placental tissues and explants were embedded in Tissue-Tek® Optimum cutting temperature (OCT) slurry (4583, VWR) and flash frozen using a dry ice-isopentane slurry. Samples were stored at −80°C until further processing.

### Immunostaining and visualisation

Whole-mount placental explant biopsies (0.5cm^3^) were fixed in 4% paraformaldehyde in PBS for 45 minutes at room temperature. Placental explants were permeabilized and blocked with 2% Triton-X-100 with 10% FBS at room temperature for 1 hr. Placental tissue sections from OCT emendation were blocked with 4% paraformaldehyde for 30 minutes at room temperature. Whole-mount and tissue sections were stained with primary antibodies (Supplementary table 2) in 0.02% Triton-X-100 in 2% FBS overnight at 4 °C. Matched secondary antibodies (Supplementary table 2) species (if using unconjugated primary) were diluted (Supplementary table 2) in 0.02% Triton-X-100 and incubated at room temperature for 3 hr. Whole mount and sections were washed three times with 1x PBS. DAPI (4’,6-Diamidino-2-Phenylindole, Dilactate) nuclear dye (564907, BD) was diluted to a concentration of 2μg/ml in PBS and added for 30 minutes before imaging. Whole mount and sections were washed three times with 1x PBS. Whole mounts were mounted in the µ-Dish 35 mm chamber system (81156, Ibidi). Tissue sections were air-dried and mounted using ProLong™ Gold Antifade Mountant (P36930, Invitrogen). Imaging was performed using the Multiphoton DLS SP8 confocal microscope (Leica Microsystem) and process using LAS-X Application Suite X Software (Leica Microsystem) and ImageJ v1.52.

### Multiplexed smFISH and high-resolution imaging

Large tissue section staining and fluorescent imaging were conducted largely as described previously^99^. Sections were cut from fixed frozen samples embedded in OCT at a thickness of 10 or 11 μm using a cryostat, placed onto SuperFrost Plus slides (VWR) and stored at −80 °C until stained. Tissue sections were then processed using a Leica BOND RX to automate staining with the RNAscope Multiplex Fluorescent Reagent Kit v2 Assay (Advanced Cell Diagnostics, Bio-Techne), according to the manufacturers’ instructions. Probes are listed in Supplementary table 3. Before staining, human fresh frozen sections were post-fixed in 4% paraformaldehyde in PBS for 15 minutes at 4 °C, then dehydrated through a series of 50, 70, 100 and 100% ethanol, for 5 minutes each. Following manual pretreatment, automated processing included epitope retrieval by protease digestion with Protease III for 10 minutes before probe hybridization. Tyramide signal amplification with Opal 520, Opal 570 and Opal 650 (Akoya Biosciences) and TSA-biotin (TSA Plus Biotin Kit, Perkin Elmer) and streptavidin-conjugated Atto 425 (Sigma Aldrich) was used to develop RNAscope probe channels.

Stained sections were imaged with a Perkin Elmer Opera Phenix High-Content Screening System, in confocal mode with 1 μm z-step size, using a ×20 (numerical aperture (NA) 0.16, 0.299 μm per pixel), ×40 (NA 1.1, 0.149 μm per pixel). Channels were as follows: DAPI (excitation 375 nm, emission 435–480 nm), Atto 425 (excitation 425 nm, emission 463–501 nm), Opal 520 (excitation 488 nm, emission 500–550 nm), Opal 570 (excitation 561 nm, emission 570–630 nm) and Opal 650 (excitation 640 nm, emission 650–760 nm).

### Luminex assay

Luminex assays were performed using the Milliplex Human Cytokine/Chemokine Magnetic Bead Panel (HCYTOMAG-60K-41, Merck Millipore) according to the manufacturer’s instructions. Cell culture supernatants were run in duplicate wells and read on the FLEXMAP 3D instrument (Thermo Fisher). Read outs were analysed using Belysa® analysis software (Thermo Fisher) to evaluate cytokine/chemokine concentrations according to the spline curve-fitting method.

### Glucose uptake assay

Trophozoites stage parasites were washed twice with glucose-free RPMI 1640 media (11879020, Thermo Fisher). Parasites were adjusted to 3% final hematocrit in glucose-free media containing 1:10,000 dilution of glucose probe labelled with red fluorescence (UP03-10, Dojindo EU) for 15 minutes at 37°C protected from light. Unlabelled parasites were included as a control. The parasites were then washed twice with an ice-cold WI buffer (UP03-10, Dojindo EU) and immediately analysed by CytoFLEX flow cytometer (Beckman Coulter). Non-GFP *P. falciparum* 3D7-EXP1wtlow-Ty1 (sfa32) parasites were co-stained with SYBR Green prior to flow cytometry acquisition. Gating strategy is according to Supplementary Figure 11.

### Alignment and quantification of explant sc or snRNA-seq data, doublet detection and clustering

The sequencing data from 10x Genomics was aligned and quantified using STARsolo v2.7.9a^100^ against the GRCh38 (ver. 2020-A, CellRanger) genome. For *P. falciparum* infection assay, sequencing data was aligned to a combined reference of human GRCh38 and *P. falciparum* 3D7 genome v3 (https://www.sanger.ac.uk/resources/downloads/protozoa/). For *T. gondii* infection assay, a combined reference of human GRCh38 and *T. gondii* GTI genome GCA_000149715.2. Gene expression (GEX) doublet scoring was done with Scrublet^101^, Souporcell^102^ was used to flag doublets by genotype for multiplexed samples. For *T. gondii* infected explants, *T. gondii* counts were quantified per cell. Multiplexed samples were deconvoluted using Souporcell with a k=2. Cells with >100 genes expressed, >300 unique molecular identifier (UMI) counts and with mitochondrial genes <20% were kept. Nuclei with >1000 genes expressed, >2000 UMI counts and with mitochondrial genes <5% were kept. Souporcell (k=2, and k=4) was used to deconvolve the Maternal and fetal origin of nuclei/cells. To estimate the cell cycle phase of each cell (G1, S or G2/M), we used the expression of G2/M and S phase markers and classified the barcodes using the method implemented in Scanpy *score_genes_cell_cycle* function. Principal component analysis (PCA) was used for dimensionality reduction, neighbour identification and Leiden clustering were run. Cells not expressing human genes, and with high expression of *Toxoplasma* markers were flagged as Tg tachyzoites and excluded from the downstream analysis. The samples were integrated using scVI v0.10.1^103^ (batch= donor+sample, raw counts) using 30 latent variables. Leiden clustering and uniform manifold approximation and projection (UMAP) were used for visualisation.

### Explant scRNA-seq and snRNA-seq annotation and analysis

The cells were annotated using the predictions of a logistic regression model built with the 3000 top highly-variable genes identified using *highly_variable_genes* SeuratV3^104^ from Scanpy, and our prior knowledge of gene markers. The model was built using scaled log-normalised counts for training and test sets, max_iter = 5000. Clusters with a high doublet score or high proportion of donor doublets tagged by Souporcell were excluded. Low QC clusters with low gene counts, high expression of apoptosis markers, and no cluster-specific genes, were removed as well. Zoom in steps in all major cell compartments were performed, where the steps mentioned above were repeated in each compartment to increase granularity. DEG analysis was done on log-normalised counts using the one-sided, non-parametric Wilcoxon Rank Sum test implemented in the *FindMarkers* function with Seurat v.3.2.2 between infected vs uninfected samples. Only those genes showing a change in the same direction in all donors were kept for further analysis. Gene Ontology (GO) enrichment analysis was done using the differentially expressed genes as input for Metascape^105^. Ligand-receptor interactions of secreted cytokines were inferred using the repository of curated ligand-receptor interactions CellPhoneDB ^106^. Cells from *T. gondii-*infected samples with a higher proportion of parasite counts than the expected by chance in uninfected samples were annotated as having intracellular parasites, “*Tg-intracellular*”. Those cells from infected samples which did not fulfil the threshold were tagged as “*No-intracellular Tg*”.

### Explant hashing scRNA-seq analysis

For hashing samples, Both libraries from hashing and GEX samples were processed and aligned together with Cellranger v3.1. Antibody capture/HTO reads with fewer than 500 unique molecular identifier counts (UMI) and doubles were excluded. Cell barcodes and hash barcodes were merged to create a GEX matrix with only overlapping barcodes, with each cell containing the HTO identifier. The GEX matrix was then processed according to Supplementary figure 3.

### *P. falciparum* scRNA-seq analysis

For malaria parasite reads, low-quality cells with <200 genes and <300 UMI counts were removed. Reads mapping to humans and parasites were identified. Total read counts per cell mapping to >90% of *P. falciparum* genome was included in the analysis. The samples were integrated using *harmonypy* v0.0.9^107^ (batch=parasite batch). Leiden clustering and UMAP were used for visualisation and cell cluster were annotated based on prior knowledge of intra-erythrocytic life cycle marker genes and Term Frequency-inverse Document Frequency (TF-IDF) ordering for top marker genes for each Leiden cluster. DEG analysis was performed using the one-sided Wilcoxon Rank Sum test implemented in the *FindMarkers* function with Seurat v.3.2.2 comparing between ‘pf_b’ vs ‘pf_iv’ samples.

## Acknowledgements

This publication is part of the Human Cell Atlas. We thank Pascale Cossart (Pasteur Institute, Paris) for supplying the *L. monocytogenes* EGD-cGFP strain. We thank Sophie Adjalley for the PfNF54eGFP strain. We thank Tobias Spielmann (Bernhard Nocht Institute for Tropical Medicine) for supplying the 3D7-EXP1wt^low^-Ty1 (*sfa32*) condΔEXP1 line. We gratefully acknowledge the Sanger Cellular Generation and Phenotyping (CGaP) Core Facility and the Sanger Core Sequencing pipeline for support with sample processing and sequencing library preparation; Bee Ling and the Cytometry Core facility for their support and training; Tarryn Porter, Yvette Wood, Di Zhou and the Cellular Genetics Wet Lab Support Team for their help in the wet lab logistics; David Goulding and the Advanced Light and Electron Microscopy suite; Nick Thomson and Sally Kay for bacteria culture facilities. We are grateful to Maximiliano Gutierrez and Amanda Oliver for their advice in metabolic dynamics, and to Aidan Maartens for proofreading. Placental material was provided by the Joint MRC-Human Cell Atlas (MR/S036350/1). The authors are grateful to patients for donating tissue for research.

## Data availability

Data generated in this study will be deposited in EMBL-EBI ArrayExpress. All codes used for data analysis will be available at https://github.com/ventolab.

## Supplementary Materials

### Supplementary Figures

**Supplementary figure 1. Workflow, donor representation and characterisation of human placental explants by scRNA-seq.**

**a)** Image of placental explant on a Matrigel-coated transwell at d0 (left) and d2 (right) of culture. Note the outgrowth of EVT from the d2 explant. **b)** Workflow for scRNA-seq processing of placental explants and CD45+ cells enrichment (see Methods). **c)** List of donor IDs used for fresh vs day 2 explant characterisation in figure 1b, 1c, and supplementary figure 1e. **d)** UMAPs of cell lineages from placental explants form two donors. Colours correspond to cell lineages in the schematic in Figure 1a. Bar plots underneath show proportions of cell types in 3 regions of the same placental explant (labelled with hashtag-oligos HTO5, HTO4 or HTO3). HB cells have been omitted for clarity, as they represent the majority of cells captured. **e)** Dot plots showing variance-scaled, log-transformed expression in selected cell type lineage clusters. **f)** List of donor IDs used for placental infection assay with all three pathogens for scRNA-seq, snRNA-seq and Luminex assay.

**Supplementary figure 2. Characterisation of human placental explants by immunofluorescence, flow cytometry and scRNA-seq.**

**a)** Representative immunofluorescence staining of d0 placenta, stained for SDC1 (SCT marker, magenta) and CD54 (macrophage marker, green), co-stained with DAPI (blue). Image tracings of microscopic images indicate the SCT and VCT areas of the imaged placental villi. **b)** Immunofluorescence staining of placental explants after 2 days in culture (d2), stained for SDC1 (SCT marker, magenta) and CD54 (macrophage marker, green), co-stained with DAPI (blue). Image tracings of microscopic images indicate the SCT and VCT areas of the imaged placental villi. **c)** Immunofluorescence staining of placental explants after 2 days in culture (d2), stained for vimentin (VIM; fibroblast marker, magenta) and HLA-G (EVT marker, green), co-stained with DAPI (blue). Scale bars: a’), b’), c’) - 100µm, a”), b”), c”) - 50µm. **d)** FACS plots of day 0 - day 6 placental samples, together with an unstained control, stained for cellular death and CD45-positivity **e)** Barplot representing numbers of up-regulated and down-regulated differentially expressed genes (DEGs) between infected and uninfected explants per cell type lineage.

**Supplementary figure 3. Overview of analysis and quality control in scRNA-seq and snRNA-seq data for human placental explant.**

**a)** sc/snRNA-seq analysis workflow for human placental explants. **b)** UMAP of scRNA-seq datasets labelled by donor, sample ID, cell cycle and developmental stage. **c)** UMAP of snRNA-seq datasets labelled by donor, sample ID and cell cycle. **d)** Logistic regression probabilities and cell type predictions. Dot plots showing variance-scaled, log-transformed expression of known lineage markers for **e)** scRNA-seq and **f)** snRNA-seq.

**Supplementary figure 4. *In vitro* panning and qRT-PCR validation for 2D9 PfN54-eGFP.**

**a)** Left: *In vitro* panning strategy for PfN54-eGFP. Right: Related to figure 2, snRNA-seq UMAP of log-transformed, normalized expression of CSA-associated genes in placental explants. **b)** Quantitative Real-time PCR expression of panned 2D9-Pn, where n represents panning cycle number. Barplot represents log2 fold change relative to bulk unplanned PfNF54eGPF. **c)** Flow cytometry gating from single-cell malaria parasites PfNF54(non-GFP) and 2D9 PfNF54-eGFP-P10 co-stained with mitotracker.

**Supplementary figure 5. Differentially expressed genes between infected and uninfected placental explants from scRNA-seq trophoblast subclusters.**

Volcano plot showing the log2 fold change of gene expression and the statistical significance of the DEG analysis between infected and uninfected placental explants from scRNA-seq manifold. Analysis was done by cell type cluster ‘VCT_fusing’ and ‘VCT’ for *P. falciparum, L. monocytogenes* and *T. gondii*. Red dots represent DEGs with a log2 fold change of >0.1 and an adjusted *p* value of <0.05, Wilcoxon Rank Sum test.

**Supplementary figure 6. Differentially expressed genes analysis between infected and uninfected placental explants from snRNA-seq trophoblast subclusters.**

Volcano plot showing the log2 fold change of gene expression and the statistical significance of the DEG analysis between infected and uninfected placental explants from snRNA-seq manifold. Analysis was done by cell type cluster ‘SCT’, ‘VCT_fusing’ and ‘VCT’ for *P. falciparum* and *T. gondii.* Red dots represent DEGs with a log2 fold change of >0.25 and an adjusted *p* value of <0.01, Wilcoxon Rank Sum test.

**Supplementary figure 7. Gene Ontology enrichment analysis from scRNA-seq manifold.**

Bar graphs represent selected Gene Ontology (GO) terms of the upregulated (yellow) and down-regulated (grey) genes for cell type cluster ‘VCT_fusing’ and ‘VCT’ for *P. falciparum, L. monocytogenes* and *T. gondii* from scRNA-seq manifold. Bar graphs are plotted by *p*-values.

**Supplementary figure 8. Gene Ontology enrichment analysis from snRNA-seq manifold.**

Bar graphs represent selected Gene Ontology (GO) terms of the upregulated (yellow) and down-regulated (grey) genes for cell type cluster ‘SCT’, ‘VCT_fusing’ and ‘VCT’ for *P. falciparum* and *L. monocytogenes* from snRNA-seq manifold. Bar graphs are plotted by *p*-values.

**Supplementary figure 9. Flow cytometry profiling for 2D9 PfNF54-eGFP.**

**a)** Representative flow cytometry gating from single-cell population of 2D9 PfNF54-eGFP ‘pf_iv’, ‘pf_b’ and ‘pf_nb’ fraction, co-stained with mitotracker. **b)** Bar plot representing average fold change of total parasite numbers relative to ‘pf_iv’. Asterisk * significance at *p* <0.05 using non-parametric Mann-Whitney test, n=3.

**Supplementary figure 10. Overview of analysis and quality control in scRNA-seq for P. falciparum.**

**a)** scRNA-seq analysis workflow for *P. falciparum*. **b)** UMAP of *P. falciparum* scRNA-seq datasets labelled by percentage of read counts mapping to *P. falciparum*, parasite culture batch and samples. **c)** Dot plots showing variance-scaled, log-transformed expression of known intraerythrocytic developmental cycle genes in the *P. falciparum* scRNA-seq datasets.

**d)** UMAP of log-transformed, normalized expression for *VAR2CSA*. Volcano plot showing the log2 fold change of gene expression and the statistical significance of DEG analysis between ‘pf_b’ and ‘pf_iv’ for **e)** early trophozoites and **f)** late trophozoites. Red dots represent DEGs with a log2 fold change of >0.25 and an adjusted *p* <10e-30.

**Supplementary figure 11. Glucose uptake assay for *P. falciparum* 3D7-EXP1wt^low^-Ty1\** Representative flow cytometry gating from single-cell population of *P. falciparum* 3D7-EXP1wt^low^-Ty1 (*sfa32*), stained with SYBR Green (DNA) and with and without PE-labelled glucose treatment. Rapalog was added to trigger the DiCre-induced excision of *exp1*. Uninfected erythrocytes were included as a negative control. Single-cell population of *P. falciparum* was gated for positive SYBR Green signal to detect total infected erythrocytes. Total infected erythrocytes were then gated for positive PE signal (glucose uptake). To account for parasite stage transition from early to late trophozoites, parasites were gated by SYBRlow (early) and SYBRhigh (late) intensity. The proportion of PE stained parasites were evaluated within the SYBRlow and SYBRhigh population.

### Supplementary Tables

**Supplementary table 1**

List of primers used for quantitative Real-Time PCR.

**Supplementary table 2**

List of primary and secondary antibodies used for immunofluorescence.

**Supplementary table 3**

List of probes used for multiplexed smFISH and high-resolution imaging.

