## Supplementary Table 1 for "Early infection response of the first trimester human placenta at single-cell scale"

| Gene_product | GeneID | Forward (5'->3') | Reverse (5'->3') |
|---|---|---|---|
| var2csa | PF3D7_1200600 | TGGTGATGGTACTGCTGGAT | TTTATTTTCGGCAGCATTTG |
| upsA | PF3D7_0400400 | ATATGGGAAGGGATGCTCTG | TGAACCATCGAAGGAATTGA |
| upsB | PF3D7_0100100 | TGCGCTGATAACTCACAACA | AGGGGTTCATCGTCATCTTC |
| upsC | PF3D7_0421100 | TGGCGTCCACTCCGACA | GAGCCTTCGGATTCATCGTG |
| arginine-tRNA ligase | PF3D7_1218600 | AAGAGATGCATGTTGGTC | GTACCCCAATCACCTACA |
