## Supplementary Table 2 for "Early infection response of the first trimester human placenta at single-cell scale"

List of primary antibodies used:

| Name | Brand | Catalog # | Clone | Dilution |
|---|---|---|---|---|
| CD138 (Syndecan-1) FITC Antibody, anti-human, REAfinity | Miltenyi Biotec | 130-115-478 | REA929 | 1:100 |
| Anti-Green fluorescence protein (GFP) | Abcam | ab13970 | NA | 1:200 |
| Anti-Listeria monocytogenes | Abcam | ab35132 | NA | 1:50 |
| Anti-Toxoplasma gondii antibody | Abcam | ab23507 | NA | 1:1000 |
| CD138 (SCD1) PE Antibody | Miltenyi Biotec | 130-119-840 | 44F9 | 1:200 |
| Alexa Fluor® 488 anti-human CD64 Antibody | Biolegend | 305010 | 10.1 | 1:200 |
| Recombinant Anti-Vimentin antibody | abcam | ab92547 | EPR3776 | 1:350 |
| Anti-HLA G antibody [MEM-G/9] PE | abcam | ab24384 | MEM-G/9 | 1:25 |

List of secondary antibodies used:

| Name | Brand | Catalog # | Clone | Dilution |
|---|---|---|---|---|
| Goat anti-Rabbit IgG (H+L) Cross-Adsorbed Secondary Antibody, Alexa Fluor™ 488 | Invitrogen | A11008 | NA | 1:600 |
| Donkey anti-Goat IgG (H+L) Cross-Adsorbed Secondary Antibody, Alexa Fluor™ 647 | Invitrogen | A21447 | NA | 1:600 |
| Donkey anti-Rabbit IgG (H+L) Highly Cross-Adsorbed Secondary Antibody, Alexa Fluor™ Plus 555 | Invitrogen | A32794 | NA | 1:600 |
