## Supplementary Table 3 for "Early infection response of the first trimester human placenta at single-cell scale"

| **Probe** | **Catalogue number** |
|---|---|
| CCL4 | 455348 |
| Hs-CCL20-C2 | 409618-C2 |
| Hs-IL1B-C4 | 310368-C4 |
| Hs-CXCL3-C4 | 1002158-C4 |
| Hs-PTGS2-O1-C2 | 508-118-C2 |
| Hs-GATA3-C3 | 403558-C3 |
| Hs-MMP1 | 412648 |
